# Activity-dependent glassy cell mechanics I : Mechanical properties measured with active microrheology

**DOI:** 10.1101/2022.09.02.506288

**Authors:** H. Ebata, K. Umeda, K. Nishizawa, W. Nagao, S. Inokuchi, Y. Sugino, T. Miyamoto, D. Mizuno

## Abstract

Active microrheology was conducted in living cells by applying an optical-trapping force to vigorously-fluctuating tracer beads with feedback-tracking technology. The complex shear viscoelastic modulus *G*(*ω*) = *G*′(*ω*) – *iG*″(*ω*) was measured in HeLa cells in an epithelial-like confluent monolayer. We found that *G*(*ω*) ∝ (−*iω*)^1/2^ over a wide range of frequencies (1 Hz < *ω*/2*π*<10 kHz). Actin disruption and cell-cycle progression from G1 to S and G2 phases only had a limited effect on *G*(*ω*) in living cells. On the other hand, *G* (*ω*) was found to be dependent on cell metabolism; ATP-depleted cells showed an increased elastic modulus *G*′(*ω*) at low frequencies, giving rise to a constant plateau such that *G*(*ω*) = *G*_0_ + *A*(−*iω*)^1/2^. Both the plateau and the additional frequency dependency ∝ (−*iω*)^1/2^ of ATP-depleted cells are consistent with a rheological response typical of colloidal jamming. On the other hand, the plateau *G*_0_ disappeared in ordinary metabolically active cells, implying that living cells fluidize their internal states such that they approach the critical jamming point.

**Statement of Significance:** Intracellular mechanical properties were measured using optical-trap-based microrheology. Despite expectations to the contrary, shear viscoelasticity was hardly affected by reorganization of cytoskeletal structures during cell-cycle progression (G1 to S and G2 phases), nor by artificial disruption of the actin cytoskeleton induced by chemical inhibitors. Rather, the mechanics of cell interiors is governed by the glassy cytoplasm. Cells depleted of ATP solidified, whereas living cells that maintained metabolic activities were more fluid-like. Instead of a completely fluid response, however, we observed a characteristic power-law viscoelasticity *G*(*ω*) ∝ (−*iω*)^1/2^ over the whole range of frequencies measured. Based on our current understanding of jamming rheology, we discuss how cells fluidize their internal state in a way that pushes the system towards the critical jamming transition.

## INTRODUCTION

The mechanics of cells is a fundamental aspect of biology; it governs the dynamics of functional bio-macromolecules that perform various physiological processes (1–3). For instance, motor proteins and other mechanoenzymes change their shapes to catalyze biochemical reactions (4, 5). They also need to be present at the right place at the right time (1); thus their performance depends on the mechanical properties (*i.e*., fluidity) of the surrounding medium. At the same time, mechanical stability is also necessary for cells in order for them to maintain their complex internal organization. Living cells thus meet these apparently conflicting requirements: dynamic fluidity and mechanical stability. However, the physical machinery underlying the regulation of intracellular mechanics remains elusive (6, 7).

It has been believed that the mechanics of cells is determined by the properties of the cytoskeleton, a network of interlinking protein filaments (8, 9). After decades of intense research, a consensus has been reached that the mechanical properties of the cytoskeleton is described using a theory based on the semiflexible polymer network model (10). However, in living cells, the interstices of the sparse network of cytoskeletons are densely filled with other colloidal constituents such as proteins, nucleic acids, polysaccharides, and their assemblies such as ribosomes (2, 11–13). In contrast to the cytoskeleton, it remains to be elucidated whether and/or how the cytoplasm excluding the cytoskeleton (hereafter referred to as just the cytoplasm) contributes to intracellular mechanics. As it was recently pointed out, cytoplasmic rheology may be similar to that of dense colloidal suspensions close to jamming (3, 7).

As is typical in soft materials, both the cytoskeleton and cytoplasm respond in a highly nonlinear manner to mechanical perturbations. For instance, when subjected to mechanical loads such as shear stress or a locally applied force, semiflexible polymer networks typically stiffen (14–20) while dense colloidal suspensions are fluidized (21–24). Active mechanoenzymes such as molecular motors also apply forces to cytoskeletons (19) which will eventually be transmitted to the surrounding cytoplasm. Conformational changes of enzymes that occur during enzymatic reactions (4, 5) may also mechanically perturb a crowded environment. It is then reasonable to expect that driving the cell interior out of equilibrium will lead to an emergence of a complex intracellular mechanics.

Living cells dynamically change their internal organization during their cell-cycle progression. They replicate chromosomes, double cytoplasmic components and reconstruct cytoskeleton structures (25). Accordingly, it has been reported that the elastic and viscous moduli of cells depend on the cell-cycle phase (26–28). All the factors listed above, *i.e*., the cytoskeleton, cytoplasm (in the interstices of cytoskeletons), their reorganization during cell-cycle progression, and their metabolic activity, seem to be integrated into intracellular mechanics. Prior studies of intracellular mechanics mostly focused on the cytoskeleton, but attention has rarely been paid to the cytoplasmic contribution. In this study, we aim to investigate how the mechanics of the cytoplasm is regulated by metabolic activities, using interphase cells which make up majority of our body and also cell cultures.

Microrheology (MR) is a method to investigate the viscoelastic properties of a medium at microscopic (~ μm) length scales (29), by tracking the motion of micron-sized tracer beads imbedded in a specimen (30, 31). In active microrheology (AMR) (32, 33), an external force is applied to a tracer bead, and its displacement response is measured. When a sinusoidal force is applied to the bead, we obtain the complex shear viscoelastic modulus *G*(*ω*) = *G*′(*ω*) – *iG*″(*ω*) of the surrounding medium as a function of the angular frequency *ω*. Here, *G*’ and *G*” are the real and imaginary part of the complex modulus *G*(*ω*). AMR can be performed simultaneously by applying an optical-trapping force to the probe particle, and measuring the probe displacements using the back-focal-plane laser interferometry technique (BFPI) (34). Because of the vigorous non-thermal fluctuations, a probe particle in living cells moves out of the laser focus within the experimental time scale. Although a strong laser could trap the probe in the laser focus, it also hinders bead motion, causing large errors in estimating the intrinsic response and fluctuations that should have occurred in the absence of optical trapping. In a prior study, we therefore developed an optical-trap-based MR implemented with a three-dimensional feedback-controlled sample stage (feedback-tracking MR, see supplementary S1) (6, 35). This technique allowed us to stably track a vigorously-fluctuating probe particle in living cells with a laser power as small as 0.4 ~ 1.8 mW, significantly lower than ordinary optical-trapping measurements (33, 36). As seen in Fig. S2, this non-invasive technology is crucial for conducting optical-trap-based MR in living cells.

Using this MR technique, we measured the intracellular mechanics of living HeLa cells. To focus on the cytoplasmic mechanics, the contribution of the cytoskeleton was reduced by (1) using probe particles that do not bind to cytoskeletal filaments, and (2) measuring the location where actin cytoskeletons are supposed to be sparse, *i.e*., the middle of cells cultured in an epithelial-like monolayer sheet. We found that the cell-cycle progression during interphase and cytoskeletal organization merely have a marginal effect on intracellular mechanics measured in this way. The complex shear viscoelastic modulus *G*(*ω*) did not show large variation from cell to cell; they consistently showed a frequency dependence predicted for disordered colloidal jamming *G*(*ω*) = *A*(−*iω*)^1/2^ (6, 7).

On the other hand, the intracellular *G*(*ω*) was highly dependent on metabolic activities. The viscoelasticity was seen to increase in ATP-depleted HeLa cells. A plateau elasticity *G*_0_ appeared which is real and constant over the frequency range, *i.e*., *G*(*ω*) = *G*_0_ + *A*(−*iω*)^1/2^. We discuss how these observations are consistent with what is expected for a glassy cytoplasm (3, 7, 37). Although it is inferred that the metabolic activity fluidized the cytoplasm (3, 7), a typical fluid-like response *G*(*ω*) ∝ –*iωη* was not observed in the range of frequencies measured (0.1 Hz ~ 10^4^ Hz). Instead, our results imply that the rheological properties typical for critical jamming (38, 39) is achieved during the active fluidization of the glassy cytoplasm in living cells (40).

## MATERIALS AND METHODS

### Cell preparation

HeLa cells and HeLa/Fucci2 cells (Riken, Cell Bank) were seeded on fibronectin-coated glass-bottom petri dishes in Dulbecco’s modified Eagle’s medium (Wako, D-Mem, high glucose) with glucose (1 mg/ml), penicillin (100 U/ml), streptomycin (0.1 mg/ml), amphotericin B (250 mg/ml), and 10% fetal bovine serum (FBS) at 37°C. Cells were cultured in a CO_2_ incubator until they formed a confluent epithelial-like monolayer sheet. The surface of the probe particles (melamine particles, 1 μm diameter, micro Particles GmbH) were coated with polyethylene glycol (PEG) strands (NANOCS, mPEG-NH2, 1000Da, PG1-AM-1k) (35), PEG coating generally passivates probe surfaces; in aqueous environments, hydrophilic PEG acts as a polymer brush and prevents sticking to other objects or to other molecules. Probe particles were introduced into cells using a gene gun (Bio-Rad, PDS-1000/He). Excess beads that did not enter the cells were removed by washing the dishes with phosphate-buffered saline. After replenishing the dishes with fresh culture medium, they were placed in an CO_2_ incubator at least overnight to allow for the recovery from cell damage. Since supplying CO_2_ causes additional noise, MR experiments were performed in a CO_2_ -independent culture medium (Gibco, L-15 medium) with 10% FBS serum. All measurements were performed at 37°C. For MR experiments under ATP depletion and actin disruption, FBS was not included in the media as stated below.

### Actin disruption

To disrupt the actin network, polymerization of actin filaments was inhibited by adding chemical agents (Cytochalasin D 50 μg/ml or Latrunculin B 10 μM). After MR experiments were performed at 37°C in L-15 medium with 10% FBS (w/o chemical agents), the culture medium was replaced with the actin disruption medium (Cytochalasin D or Latrunculin B, without FBS). Cells were then incubated at 37 °C on the MR setup for ~30 min prior to the next MR measurement so that the actin network is disrupted in cells.

### ATP depletion

The high energy molecule ATP is mainly produced in cells via two metabolic pathways: glycolytic in the cytoplasm and oxidative phosphorylation in the mitochondria. Therefore, in order to deplete intracellular ATP, cells were cultured in a nutrient-free media with 50 mM 2-deoxy-D-glucose and 10 mM sodium azide (41). 2-deoxy-D-glucose and sodium azide inhibit glycolysis and oxidative ATP production, respectively.

HeLa cells were incubated in the medium for ATP depletion (L-15 medium with 50 mM 2-deoxy-D-glucose, 10 mM sodium azide, without FBS) at 37 °C for ~10 hours prior to the MR measurements so that ATP stored in cells is consumed.

### Luminescent ATP detection assay

Intracellular ATP was measured using a CellTiter-Glo® Luminescent Cell Viability Assay kit (Promega, G7570) according to the manufacturer’s instructions. Briefly, 5,000 cells were plated on a 96-well plate and incubated for 24 h at 37°C in 5% CO_2_. Following incubation, the medium was replaced with 50 μl of either: (1) control medium [phenol-red free DMEM (Thermo Fisher, 31053028) supplemented with 4 mM glutamine (Thermo Fisher, 25030081) and 1% Penicillin-Streptomycin Solution (Sigma-Aldrich, P4333)], or (2) control medium supplemented with 50 mM 2-deoxyglucose (Nacalai, 10722-11) and 10 mM NaN3 (WAKO, 199-11095), and then incubated for 7 h at 37°C in 5% CO_2_. Subsequently, 50 μl of reagent was directly added to the well and the contents were mixed for 2 min on an orbital shaker at 25°C. Luminescence was recorded after 30 min of incubation at 25°C in the dark.

### Immunocytochemistry

HeLa cells plated on a poly-lysine-coated glass-bottom dish were fixed with 4% paraformaldehyde phosphate buffer solution (Wako, 163-20145) for 10 min at 25 °C, permeabilized with PBS containing 0.5% Triton-100 for 15 min at 25 °C, and stained with ActinRed™ 555 ReadyProbes™ Reagent (ThermoFisher, R37112) for 30 min at 25 °C. Subsequently, the cells were incubated with Hoechst 33342 (ThermoFisher, H1399) for 10 min at 25 °C for nuclear staining. The cells were washed three times with PBS after each step. Imaging was performed using an Eclipse Ti2-E microscope equipped with NIS-Elements AR imaging software. The representative images were processed by Clarify.ai and Denoise.ai using NIS-Elements AR 5.30, as previously described (42).

### AMR in living cells

The technical details of conventional optical-trapping-based AMR (without feedback operation) are given elsewhere (33). We will repeat the essence here briefly for the reader’s convenience. As shown in Figs. 1A and B, a small sinusoidal force 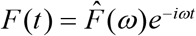 was applied to a probe particle, by optically trapping it with the drive laser (λ =1064 nm Nd:YVO_4_, Coherent Inc., Tokyo, Japan) whose focus position was controlled by an acousto-optic deflector (AOD; DTSX-400-1064, AA Opto-Electronic, Orsay, France). The probe displacement synchronous to the applied force 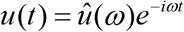 was measured using back-focal-plane laser interferometry (BFPI) conducted with a fixed probe laser (λ = 830 nm IQ1C140, Power Technology, Inc., Alexander, AR, USA) (34). Typical values of 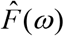 and 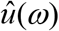 are given in Supplementary S6. The frequency response function 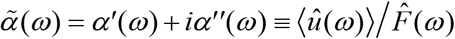 is then obtained as a complex quantity. The complex shear modulus *G*(*ω*) of the surrounding media is also obtained as a complex quantity via the Stokes relation extended to the sinusoidal response,

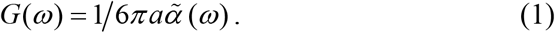

where *a* is the radius of the spherical probe particle.

**FIGURE 1.**
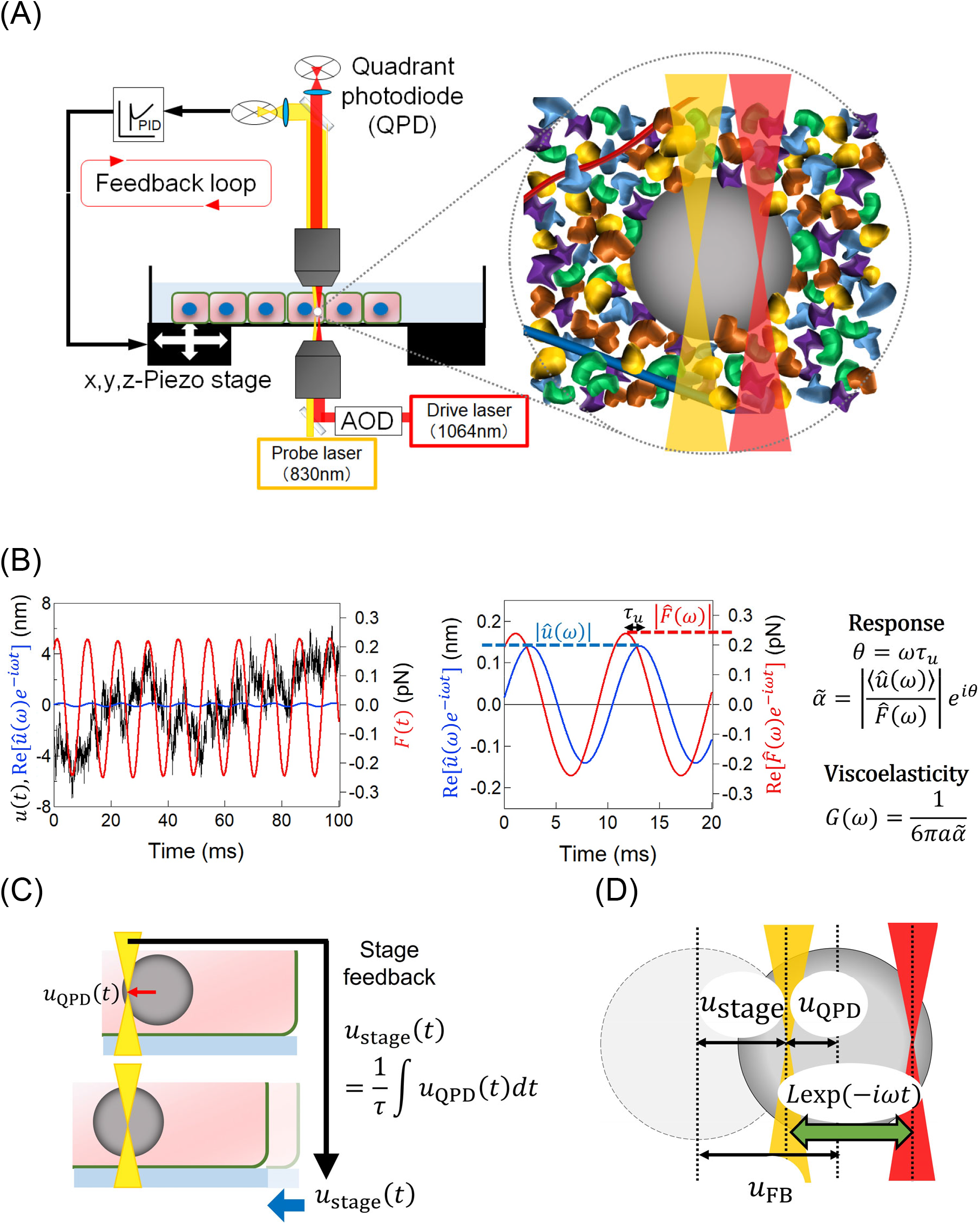
(A) Schematic of the feedback-tracking microrheology setup. A fixed probe laser is used to measure *u*_QPD_ using a quadrant photodiode (QPD). A drive laser and an acousto-optic deflector (AOD) are used to apply an oscillatory external force. The PEG-coated probe particle surrounded by dense colloidal constituents is repelled from the sparse cytoskeletal network. (B)Typical time series of the probe displacement *u*(*t*) (black line) and applied force 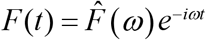 (red line). The sinusoidal signal 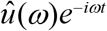 was hidden in the experimental noise (*e.g*. thermal fluctuation) but can be detected by using a lock-in amplifier. (C) Schematic illustration of stage feedback. (D) The displacements of a probe *u*(*t*) is obtained by the sum of the displacement of the piezo stage *u*_stage_(*t*) and the displacement from the focus of the probe laser. The focus of the drive laser is moved sinusoidally around the focus of the probe laser, following *L*exp(−*iωt*).

A tracer bead fluctuates vigorously in a living cell because the cytoplasm is driven far from equilibrium by the energy derived from metabolism. As explained in detail in Refs. (6, 35), such a fluctuating particle was smoothly tracked by analog PID-feedback control of the piezo-mechanical sample stage on which the specimen was placed. The optical-trapping force was then stably applied to the probe particle by the oscillating drive laser while maintaining the probe particle in the focus of the fixed probe laser. As shown in Fig. 1C, the displacement of the probe particle from the probe laser *u*_QPD_, which is measured by BFPI of the probe laser, is used to control the position of the piezo stage *u*_stage_ by finding

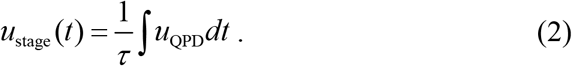

Here, *τ* (typically set at 10 ms ~ 100 ms) is the delay time for the feedback response of our experimental setup. The displacement *u*_FB_ of the probe in the coordinate system traveling with the feedback-controlled piezo stage is given as *u*_FB_ = *u*_QPD_ + *u*_stage_ (Fig. 1D). From Eq. (2), the frequency-response relation is obtained as

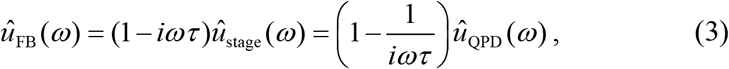

where ^ denotes the amplitude of the sinusoidal signal, *e.g*. 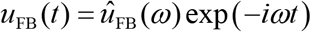. From Eq. (3), it can be seen that *u*_QPD_(*t*) and *u*_stage_(*t*) are high- and low-pass filtered from the total probe displacement *u*_FB_(*t*). Thus, the slow/large displacements of the probe are tracked by the piezo stage as *u*_stage_ whereas the fast/small displacements are measured with the QPD as *u*_QPD_. By measuring either 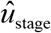 or 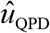 as a function of frequencies, we obtain the total probe response 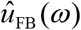 under feedback tracking. By estimating the sinusoidal force 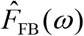 applied to the probe under the feedback (as given in Supplementary S1), we obtain the probe particle’s response function as 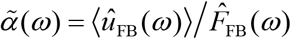. We then obtain *G*(*ω*) via Eq. (1).

### Statistical analysis

Since the distributions of *G*’ and *G*” were non-Gaussian, statistical analysis was performed by using the Mann–Whitney U test, unless otherwise stated in the figure caption. *P* values < 0.05 were considered a significant difference. All the statistical tests were performed using Igor and Matlab software. A list of all *P* values from the MR experiments is given in Supplementary S7.

## RESULTS

### Power-law intracellular rheology of untreated HeLa cells

Using feedback-tracking AMR, we measured the shear viscoelastic modulus of HeLa cells derived from a cervical cancer. Melamine particles (2*a* = 1 μm) were incorporated into cells forming an epithelial-like confluent monolayer on the surface of a glass-bottom dish. Particles incorporated at the center of cells between the cell membrane and the nuclear membrane were used as probes. The average of the complex shear modulus of the untreated HeLa cells is shown in Fig. 2 together with the standard deviation (*n* = 21). Filled circles in (A) and open circles in (B) show the real (*G*’) and imaginary (*G*”) parts of the complex shear modulus, *G*(*ω*) = *G*′(*ω*) – *iG*″(*ω*), respectively. Curves show raw data before averaging. Markers are averaged values and bars indicate the log-normal SD. Since the logarithms of mechanical properties are approximately distributed with Gaussian (41), averages and SDs were calculated for the logarithm of experimental results. Our results showed a characteristic frequency dependence of the form *G*(*ω*) ∝ (−*iω*)^1/2^. *G*’(*ω*) slightly deviates from the power-law dependency at low frequencies (≤ 1 Hz), but this may be an artifact. As shown in Supplementary S2, low-frequency fluctuations of probe particles in Madin-Darby canine kidney (MDCK) cells are decreased by prolonged irradiation with a 1.7 mW laser, implying an increased elasticity at low frequencies. Accordingly, our recent preliminary experiments performed more carefully with a reduced laser power (<0.5 mW) in MDCK cells seems to remove this deviation. This will be reported in more detail elsewhere.

**FIGURE 2.**
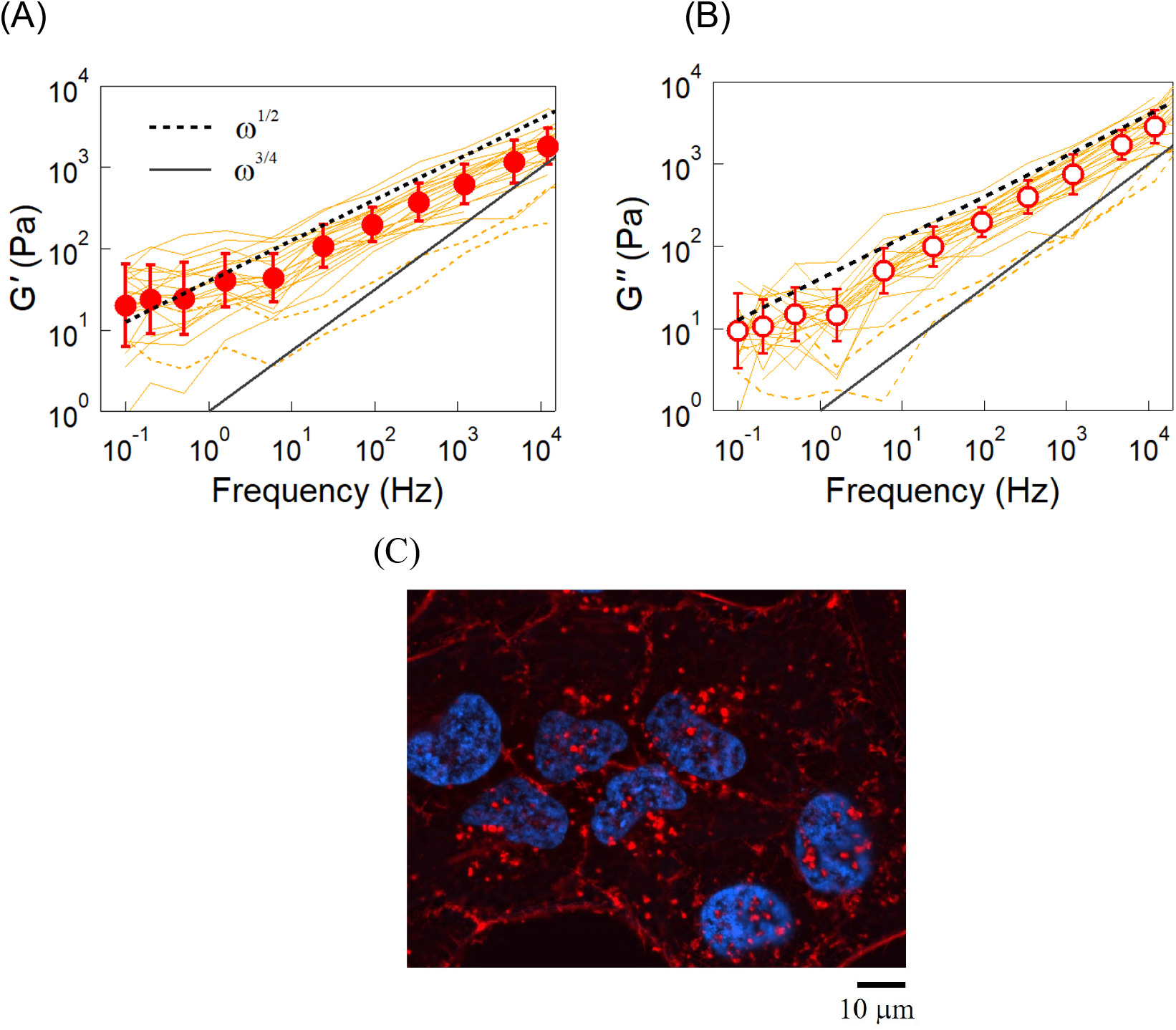
Intracellular complex shear viscoelastic modulus measured with AMR using 2*a* = 1 μm diameter melamine particles. (A) *G*’(*ω*) and (B) *G*”(*ω*) of HeLa cells in a confluent monolayer formed on a glass substrate. Curves are the raw data obtained in 23 different cells. Data shown as dotted lines were excluded from the average (n =21). Since cell cycles were not monitored, cells in different cell-cycle phases (G1, S, and G2) are included. Circles are the statistical average, and bars indicate the log-normal SD. Broken and solid lines show the power-law dependency typical of the glassy cytoplasm (∝ *ω*^1/2^) and cytoskeletal networks (∝*ω*^3/4^), respectively. The result follows ∝*ω*^1/2^ over a wide range of frequencies, indicating that the intracellular mechanics are predominantly governed by the glasssy cytoplasm. (C) Immunofluorescent images of confluent HeLa cells. The cells were stained with Rhodamine phalloidin and Hoechst33342 to visualize Factin (red) and the nucleus (blue).

It has been reported that cytoskeletal networks prepared *in vitro* exhibit an elastic plateau *G*(*ω*) ∝ (−*iω*)^0^ at low frequencies, and a power-law dependency *G*(*ω*) ∝ (−*iω*)^3/4^ at higher frequencies (16, 19, 43, 44). The frequency dependency, which is now accepted as the hallmark of cytoskeletons (20, 43), is explained by the theory for semi-flexible polymer networks (45–48). In biological cells, a *G*(*ω*) consistent with this understanding was found when MR was conducted using probe particles coupled to actin cytoskeletons (6, 41, 49). On the other hand, the power-law dependence *G*(*ω*) ∝ (−*iω*)^1/2^ observed in this study is not consistent with that expected for the cytoskeleton. It is to be noted that the probe particles used in this study were coated with polyethylene glycol polymers that generally inhibit adherence to objects in cells, including the cytoskeleton. Actin cytoskeletons are usually less expressed in confluent monolayers of cells compared to isolated cells (50). In addition, the probes were incorporated deep inside of living cells where the actin cytoskeleton is only sparsely expressed (Fig. 2C), as reported in confluent HeLa cells (51, 52) and also in confluent epithelial-like monolayers of MCF10A and MDCK (50, 53).

### Marginal contribution of the actin cytoskeleton to intracellular mechanics

In order to clarify the contribution of cytoskeletons to intracellular *G*(*ω*), we conducted AMR in HeLa cells in which F-actin was disrupted. After conducting the first AMR experiments in untreated HeLa cells, actin polymerization was inhibited by adding Cytochalasin D or Latrunculin B to the culture medium (54, 55). A second round of AMR measurements was then performed using the same particles as those used in the first AMR experiments. Since *G*(*ω*) shows a general trend proportional to (−*iω*)^1/2^, results normalized as *G*′/*ω*^1/2^ are shown in Fig. 3 in order to clarify the effect of the chemical dose at each frequency. Cytochalasin D at 50 μg/ml did not alter *G*(*ω*) in any frequency range (n = 9, Fig. 3A, and B). Statistical tests at each frequency were conducted for *G*′(*ω*) and *G*″(*ω*) before and after the addition of Cytochalasin D. P values obtained by the Mann–Whitney U test were larger than 0.05 for all frequencies measured, which indicates no significant difference. Similar results were obtained by treating cells with another inhibitor for actin polymerization, Latrunculin B at 10 μM (Fig. S3). Cells changed their morphology after the inhibitors were administrated (Fig. 3C). They became more rounded while leaving branched protrusions, lost cell-cell contact; some cells were even removed from the glass substrate. These observations indicate that the inhibitors worked effectively and disrupted cytoskeletal organization, since the actin cytoskeleton is responsible for maintaining the shape of cells and their adhesion to the substrate (56). We thus conclude that actin cytoskeletons do not contribute to the intracellular *G*(*ω*) of HeLa cells cultured in a confluent monolayer in any remarkable way.

**FIGURE 3.**
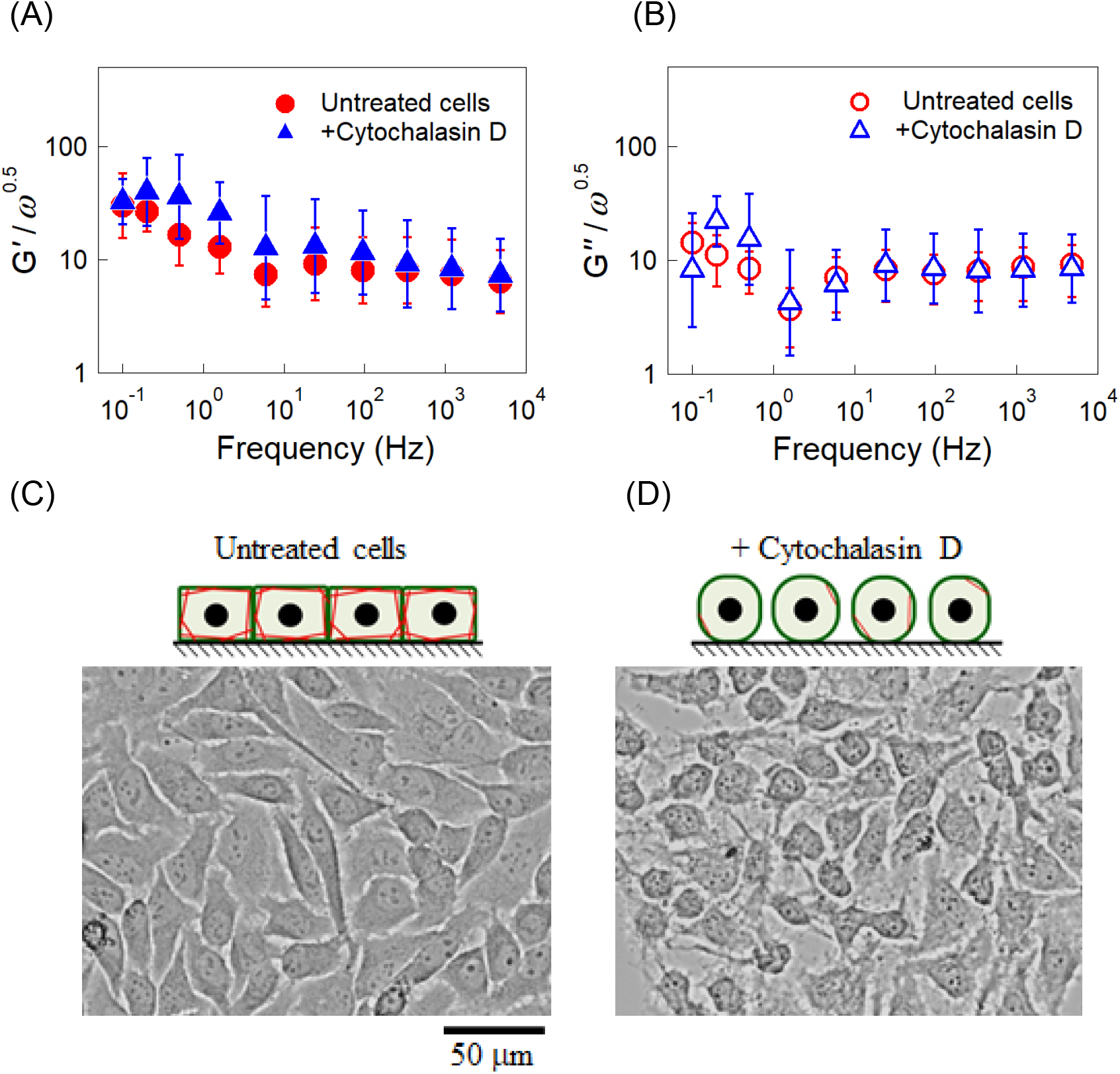
(A) *G*′/*ω*^1/2^ and (B) *G*″/*ω*^1/2^ of HeLa cells, untreated (circles) and treated (triangles) with 50 μg/ml Cytochalasin D. The same probe particles in the same cells were measured before and after adding Cytochalasin D to the culture media (n = 9). Bars indicate the log-normal SD. Similar experiments performed before and after treatment with 10 μM Latrunculin B are given in Fig. S3. (C, D) A microscopy image of HeLa cells. When actin cytoskeletons were disrupted with 50 μg/ml Cytochalasin D, cells strongly contract, leaving branched protrusions. (A, B) The Mann–Whitney U test was used to calculate the P value between untreated cells and cells treated with 50 μg/ml Cytochalasin D. No significant difference was observed for all the frequencies.

HeLa cells in confluent monolayers are laterally confined and polarized under normal conditions. They are roughly as tall as they are wide, and tightly connected to each other. In such epithelial-like tissue, actin stress fibers are densely expressed at the cell-cell and cell-substrate interfaces, and they contribute to the organization of the tissue. The disruption of actin thus remarkably affects cell morphology. In prior studies, mechanical properties of cells have been measured by using other techniques, *e.g*. AFM (atomic force microscopy) (9, 57–59), MTC (magnetic twisting cytometry) (37, 41, 60), and others (41, 61, 62). These techniques have shown that the mechanical properties of cells indeed depend on actin expression (59). It is therefore commonly believed that the mechanical properties of cells are directly correlated with their shape and cytoskeletal organization (61, 62), in contrast to our experimental results. It is to be noted, however, that the techniques used in prior studies measure the properties of cell surfaces at which actin cytoskeletons are densely expressed. Considering that F-actin is less expressed inside of the epithelial cells in a confluent monolayer, results obtained in this study are reasonable and do not contradict any of the prior measurements conducted on cell surfaces.

Cell surface elasticities measured using conventional techniques are usually widely distributed, over orders of magnitude (41, 58, 63). The organization of the actin cytoskeleton at the cell cortex is heterogeneous and changes with time. Depending on the situation, actin filaments bundle to form thick fibers, couple to cell membranes, and create tension due to actomyosin contractile activity, leading to remarkable stiffening as demonstrated **in vivo** (64–66) and **in vitro** (16, 19, 67). All these and many other factors contribute to variations in the mechanical properties of cell surfaces. On the other hand, *G*(*ω*) in living cytoplasm is found to be narrowly distributed, even though they were measured with different probes in different cells. This observation also supports the claim that the intracellular mechanics of the epithelium are not strongly associated with the actin cytoskeleton.

### Cell mechanics during cell-cycle progression (G1 to S and G2 phases)

Next, we performed feedback-tracking AMR in HeLa cells while the cell cycle progresses during interphase. HeLa cells used here (HeLa/Fucci2) were labelled using the Fucci system that enables us to identify G1 and S/G2 phases by the fluorescence color in the cell nucleus (68). In Fig. 4, we show the normalized complex shear modulus (A: *G*′/*ω*^1/2^ and B: *G*″/*ω*^1/2^) of HeLa cells as a function of frequency. Circles and squares indicate cells at G1 and S/G2 phases respectively. The intracellular *G*(*ω*) in the S/G2 phase slightly increased compared to cells at G1 phase; *G*’ and *G*” of S/G2 phase were 40% and 30% larger than those of the G1 phase on average, respectively. Note that these differences were only marginal but observed because the cell cycle phases could be resolved.

**FIGURE 4.**
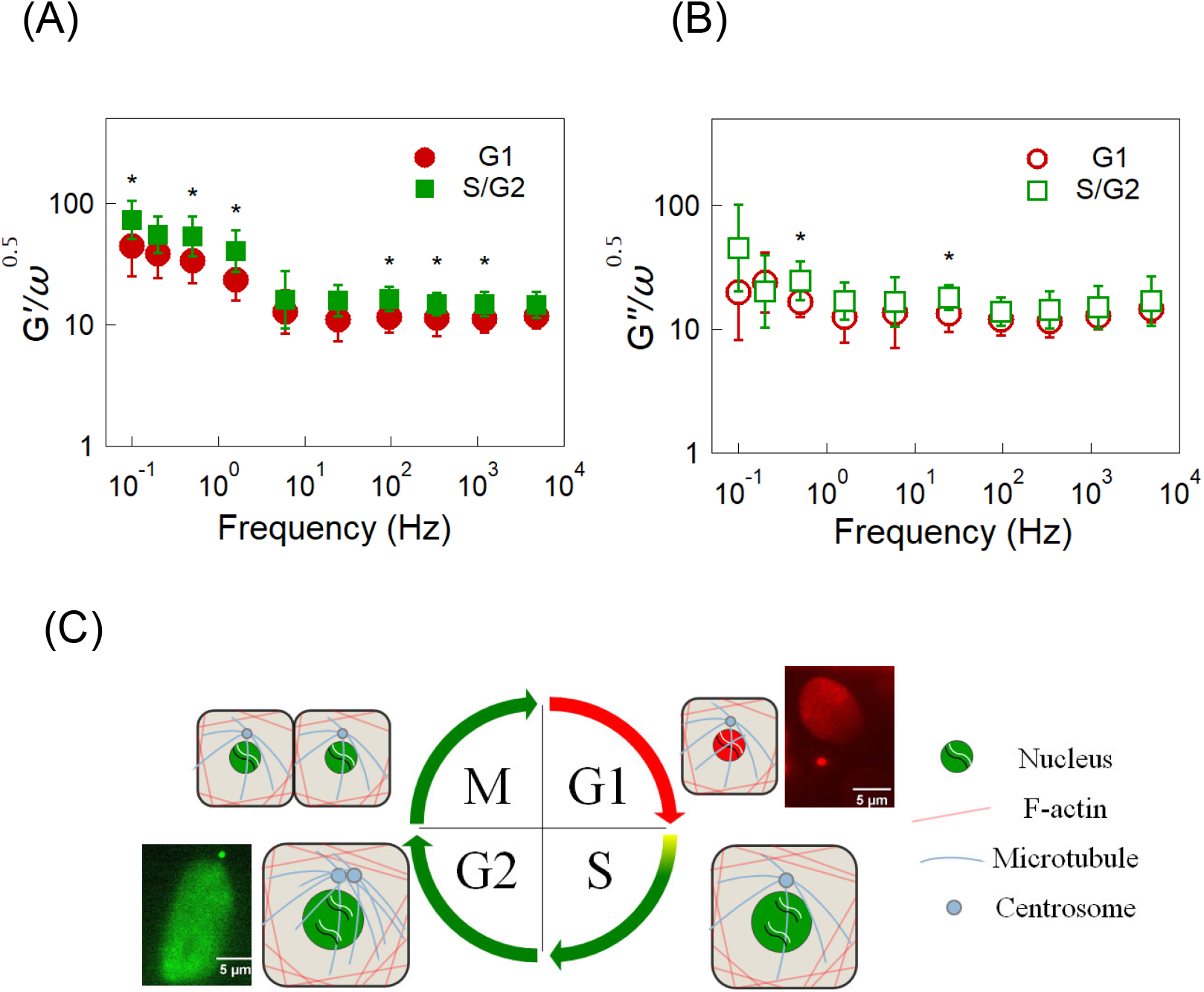
(A) *G*′/*ω*^1/2^ of HeLa/Fucci2 cells in G1 (circles, n = 12) and S/G2 phases (squares, n = 9). (B) *G*″/*ω*^1/2^ of HeLa/Fuuci2 cells in G1 (circles) and S/G2 phases (squares). Bars indicate the estimate of the log-normal SD. Remarkable differences were not observed. (C) Schematic illustration of the Fucci system used for identifying cellcycle phases. The Mann-Whitney U test was used to calculate the P value for differences between the G1 and S/G2 phases. *: P < 0.05. No symbol: P > 0.05.

In the literature, the cell-cycle dependency of cell mechanics has been investigated using atomic force microscopy (AFM) (26) and micropipette aspiration (27), both of which showed a 50~100% increase of surface elasticity of S-phase cells compared with that of G1-phase cells. Here, we emphasize that our results and prior studies do not contradict. Our AMR methodology is indifferent to F-actin, and likely measure the interstices of the sparse network of the cytoskeleton. On the other hand, prior studies measured the response of the cytoskeleton at cell surfaces whose expression is likely strongly affected by cell-cycle progression.

As explained in the introduction, various processes that occur during cell-cycle progression are inherently mechanical. Especially, cells in the M phase experience drastic variations in cytoskeletal organization (62, 69), which would affect cell mechanics derived from the cytoskeleton (28, 70, 71). When the probe particle was coupled to the cytoskeleton, MR in the M phase could exhibit mechanical properties more evidently different from other phases; the cell interior underwent softening and the viscoelastic fluid contributions increased (28). However, AMR experiments were not conducted in M phase cells in this study. Because mitosis and cytokinesis are highly dynamic processes and complete in a relatively short period of time, it was not possible to stably track the probe particle in M phase cells with the feedback technique.

### ATP-dependent intracellular mechanical properties

After finding that the actin cytoskeleton and cell-cycle progression during interphase have marginal effects on the intracellular mechanics, we now investigate the effect of metabolism on active mechanics in HeLa cells. In order to regulate the metabolic activity, cells were incubated with a culture media that causes ATP-depletion (41). Firstly, intracellular ATP was measured using the luminescent ATP detection assay. Figure 5A shows that the ATP level significantly decreased by more than an order of magnitude in ATP depleted cells. On applying AMR, it was found that the ATP-depleted cells showed an elastic plateau such that *G*(*ω*) ∝ *G_0_* + *A*(−*iω*)^1/2^ in contrast to *G*(*ω*) ∝ (−*iω*)^1/2^ for untreated cells, as shown in Fig. 5B. To quantitatively evaluate the effect of ATP-depletion on *G*(*ω*), we fitted *G*(*ω*) of untreated and ATP-depleted cells with *G_0_* + *A*(−*iω*)^1/2^ (black dashed lines in Fig. 5B). The coefficient *A* hardly changed while the elastic plateau *G*_0_ of ATP-depleted cells became ~ 10 times larger than that of untreated cells. This result clearly indicated that only *G*′(*ω*) at low frequency significantly increased due to ATP depletion. In a similar manner to previous sections, *G*′(*ω*)/*ω*^1/2^ and *G*″(*ω*)/*ω*^1/2^ of untreated HeLa cells (n = 21, red dashed line) and ATP-depleted cells (n = 9, black filled circles) are shown in Fig. 5C and D, respectively. Error bars indicate log-normal SDs except for *G*″(*ω*)/*ω*^1/2^ in ATP-depleted cells. Instead, we show the median, 25% and 75% quantiles for *G*″(*ω*)/*ω*^1/2^ in ATP-depleted cell because *G*″(*ω*) at small *ω* sometimes showed negative values due to *G*′(*ω*) ≫ *G*″(*ω*) and measurement noise. The Mann–Whitney U test for *G*’ in the low frequency range (0.1 to 94 Hz) showed a significant increase of *G*’ upon ATP depletion (P < 0.05) compared to untreated cells.

**FIGURE 5.**
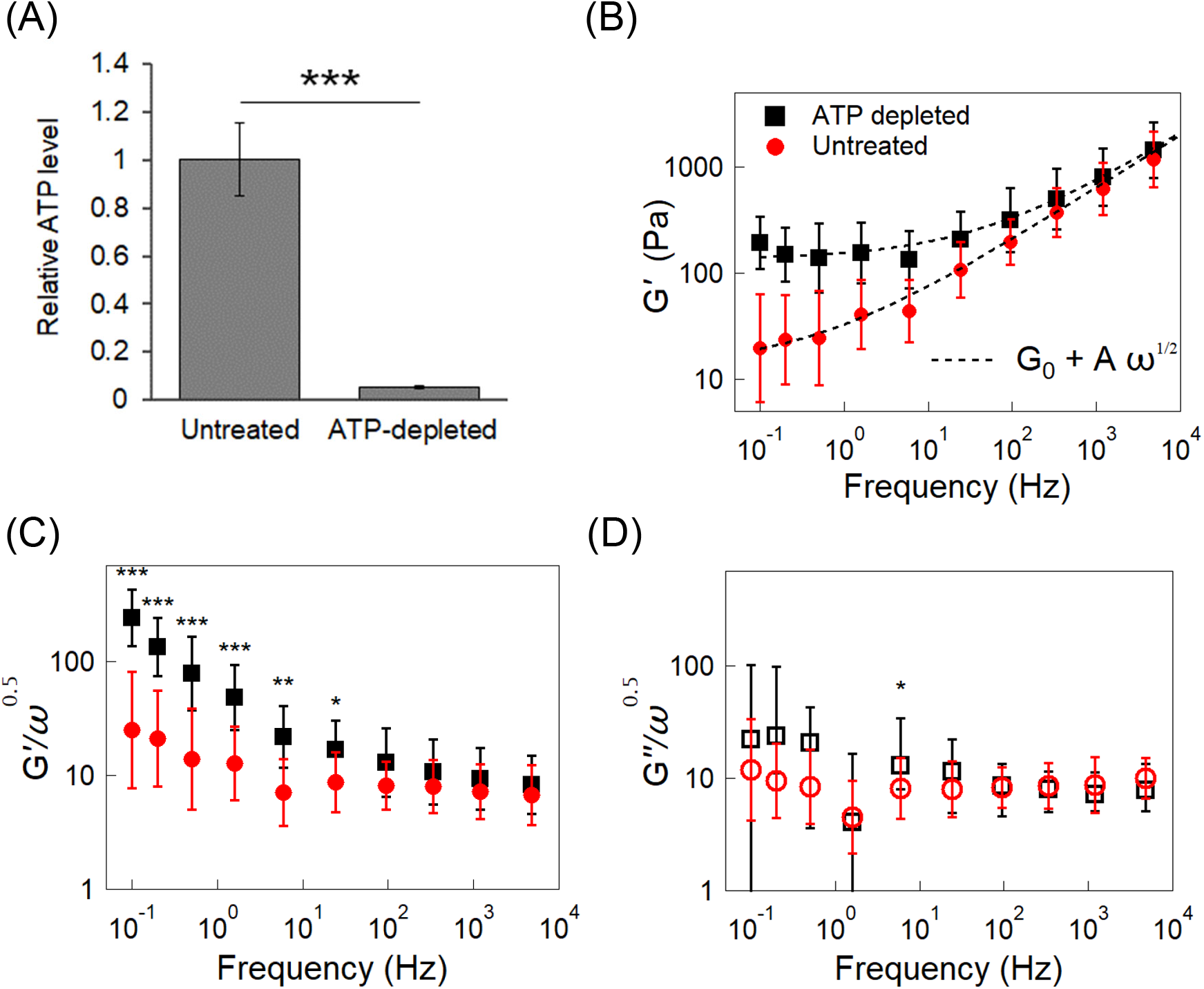
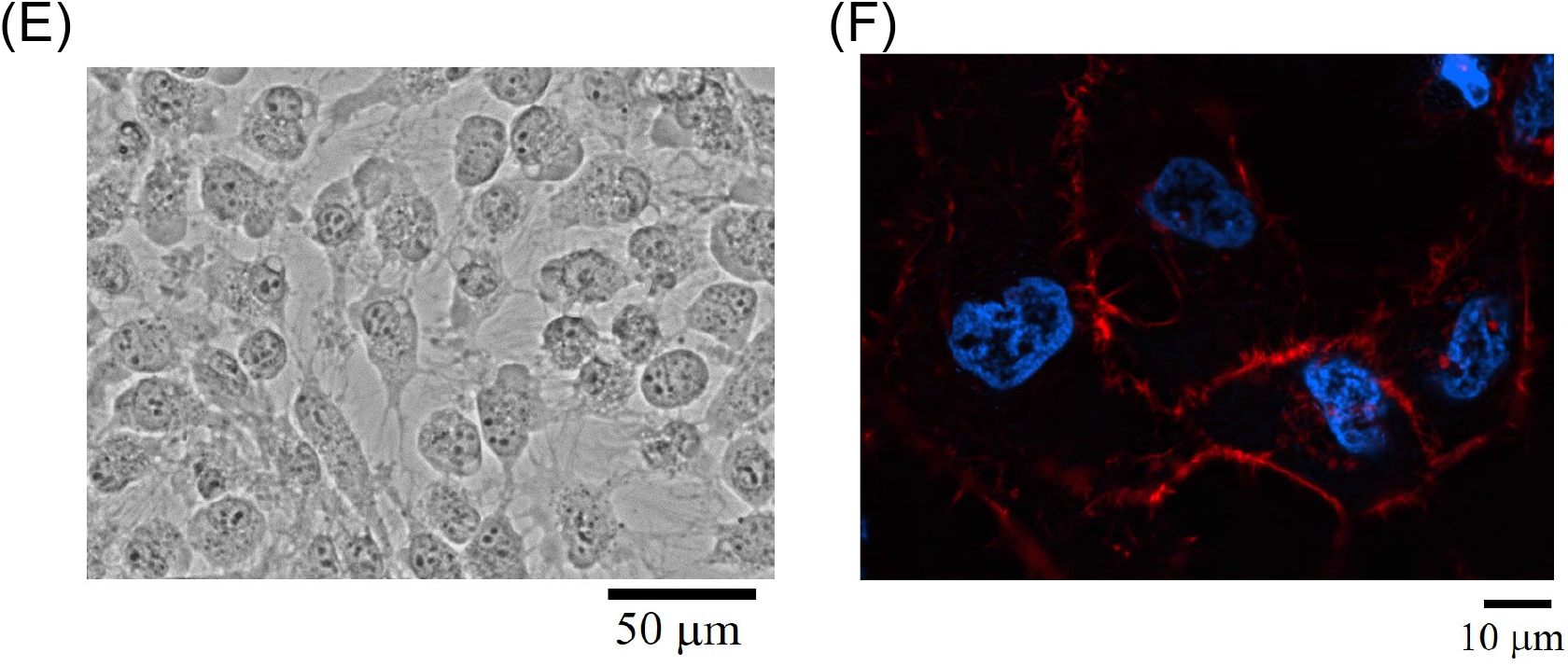
(A) Intracellular ATP level in HeLa cells treated with 50 mM 2-deoxyglucose and 10 mM NaN3 (ATP-depleted) for 7 h was measured. Quantification was performed on five independent experiments. All data are presented as mean ± standard deviation. ***: p < 0.001 (Student’s t-test). (B) *G*′(*ω*) of untreated and ATP-depleted HeLa cells. *G*(*ω*) of untreated and ATP-depleted HeLa cells are fitted by *G*_0_ + *A*(−*iω*)^1/2^. Untreated: *G_0_* = 136 Pa. *A* = 11.1 Pa s^−0.5^. ATP-depleted: *G_0_* = 13.1 Pa. *A* = 11.2 Pa s^−0.5^. (C) *G*′(*ω*)/*ω*^1/2^ and (D) *G*″(*ω*)/*ω*^1/2^ of untreated (red closed circles n = 21) and ATP-depleted HeLa cells (black open squares, n = 9). (C) Bars indicate the log-normal SD. (D) Symbols and bars of *G*″(*ω*)/*ω*^1/2^ for untreated cells indicate the log-normal mean and SD, respectively. *G*″(*ω*) of ATP-depleted cells at low frequency sometimes became negative due to *G*′(*ω*) ≫ *G*″(*ω*) and measurement noise. Thus, symbols of *G*″(*ω*)/ *ω*^1/2^ for ATP-depleted cells represent median values, and the bars connect 25% and 75 % quantiles. At low frequencies (*f* ≤ 94 Hz), *G*′(*ω*) showed a significant increase after ATP depletion (P < 0.05). (C, D) The Mann-Whitney U test was used to calculate the P value for differences between untreated and ATP depleted cells. ***: P < 0.001. **: P < 0.01. *: P < 0.05. No symbol: P > 0.05. (E) Microscope image of ATP-depleted HeLa cells. ATP-depleted HeLa cells became round and lost cell-cell contact. (F) Immunofluorescent images of ATP-depleted HeLa cells. The cells were stained with Rhodamine phalloidin and Hoechst33342 to visualize F-actin (red) and nuclei (blue).

We note that feedback-tracking MR in cells is most easily conducted when the probe particle is situated in the homogeneous part of the cell. Experimental errors increase when the probe particle is in a region close to the nucleus, cell membranes, or other large organelle, because the inhomogeneity of optical properties in such region disturbs the propagation of the probe and the drive laser. It frequently occurred that a probe particle that was used in the first AMR measurement moves to regions where measurement is no longer possible. In Fig. S4, we compare the limited number of data points (n =3) successfully measured using the same probe particle in the same cells. All the results showed a remarkable increase in viscoelasticity when ATP was depleted.

Fig. 5E shows microscope images of HeLa cells taken after ATP depletion. ATP-depleted cells lose tight adherence to the substrate and to neighboring cells. Cells change their morphology and take on more rounded shapes due to decreased cytoskeletal tension. To clarify the effect of ATP-depletion on actin cytoskeletal structures, F-actin was stained using Rhodamine phalloidin (Fig. 5F). Similar to the untreated cells (Fig. 2C), the actin cytoskeleton of ATP-depleted cells was also sparse. Thus, we concluded that the actin cytoskeleton was not responsible for the increase in the elastic plateau during ATP depletion. Note that *in vitro* cytoskeletons are remarkably stiffened under active tension or pre-stress (14–16). Consistently, the cell surface elasticity measured with AFM decreases when ATP is depleted (72, 73) likely because the active stiffening of the cytoskeleton is eased owing to the relax of actomyosin contraction at the cell cortex (61, 74–76). On the contrary, intracellular elasticity measured in this study increased significantly when the intracellular ATP was depleted. This observation again indicates that the cell cortex cytoskeleton is not relevant to intracellular mechanics. When cells are subjected to the ATP-depletion medium for a prolonged period, they could enter the path towards cell death (77, 78). The fluorescence indicator (fluorescence caspase 3) for apoptosis indeed showed positive in HeLa cells prepared using the similar protocol (Fig. S5). Except for the emergence of the elastic plateau, however, the mechanical properties typical for intracellular rheology *G*(*ω*) ∝ (*iω*)^1/2^ were still observed for ATP-depleted cells. Therefore, our results indicate that the cytoplasm with less expression of the cytoskeleton stiffens in a metabolism-dependent manner.

## DISCUSSION

It has been believed that mechanical properties of cells are mostly determined by the cytoskeleton (8, 9). This study showed, however, that intracellular mechanics in a confluent epithelium are scarcely affected by the actin cytoskeleton. Note that there are multitudes of other biomacromolecules that are crowded together in living cells. Although their effects on cell mechanics have not been fully appreciated until recently, it was shown that *in vitro* cytoplasm that lacks a cytoskeleton has glass-forming capabilities; the cytoplasm becomes glassy (*i.e*. practically solidified owing to the slowdown of dynamics) when the concentration of polymeric components is increased and/or metabolic activities are inhibited (3, 7, 37, 79). Vigorous fluctuations were observed in cells maintaining ordinary metabolism, but these were remarkably decreased in ATP-depleted cells (3, 36, 41, 80). It was then inferred that the glassy cytoplasm is fluidized by metabolic activities. However, the decrease/increase of fluctuations does not necessarily mean vitrification/fluidization. Even if glass-forming materials are deeply quenched and solidified, they could still flow if they are actively stirred (81). Besides, only observing the spontaneous fluctuations does not tell us how strongly the medium is stirred by metabolic activities. In this study, we directly measured intracellular viscoelasticity with AMR by observing a probe particle’s response to a well-determined force. Our direct measurement showed that the cytoplasm in ATP-depleted cells was glassy while cells with ordinary metabolic activities were fluidized.

Cells are constructed of soft materials that respond non-linearly to applied forces. For instance, semiflexible polymer networks (model for cytoskeletons) typically stiffen under tension (14–20) while glassy colloidal suspensions (model for the cytoplasm without the cytoskeleton) are fluidized by mechanical loads (21–24). In living cells, various mechano-enzymes such as motor proteins perform physiological functions by generating forces utilizing energy supplied by metabolism. The motor-generated forces then create non-thermal fluctuations that surpass thermal fluctuations as demonstrated by the violation of the fluctuation-dissipation theorem, both *in vivo* (6, 36, 82–85) and *in vitro* (16, 19, 33, 86, 87). Owing to the nonlinear response of constituent materials, mechanical properties of cells are profoundly altered by the active fluctuations. In the living cytoplasm, vigorous active fluctuations presumably induce structural relaxations that could not occur with thermal activation (6, 7), leading to active fluidization. Here, the fluidization of the cytoplasm under metabolic activities is consistent with the concept of a glassy cytoplasm, but not with cytoskeletons that stiffen with active metabolism (16, 19, 67).

The complex shear viscoelastic modulus *G*(*ω*) in ordinary HeLa cells showed a characteristic frequency dependence *G*(*ω*) = *A*(−*iω*)^1/2^. This dependency may seem to coincide with the prediction of the Rouse model (88) that describes the high-frequency dynamics of flexible polymers. In accordance with the theory, cross-linked poly-acrylamide gels show *G*(*ω*) ∝ (−*iω*)^1/2^, but only at high frequencies (6, 89). Networks of cross-linked semi-flexible polymers usually show *G*(*ω*) ∝ (−*iω*)^3/4^, but this changes to *G*(*ω*) ∝ (−*iω*)^1/2^ when the network is placed under tension (16, 48). Therefore, polymer networks could also show *G*(*ω*) ∝(−*iω*)^1/2^ at high frequencies when they are cross-linked. However, the polymer-network models predict that the prefactor *A* is proportional to monomer concentration (47, 88). On the other hand, *A* for extracted cytoplasm and for cytoplasm in living cells grow much-more rapidly, super-exponentially and exponentially as a function of macromolecule concentration, respectively (7). Such rapid growth of viscoelasticity has been typically observed in glass-forming materials close to the glass transition (90) but not in polymer networks. Note that the static resilience of cross-linked polymer networks should not increase at all when the metabolic activity is inhibited: in fact, the opposite should happen for crosslinked cytoskeletons (16). Also, the mechanical properties of flexible polymer networks are hardly affected by the kind of mechanical perturbations expected under physiological conditions (15, 17).

We believe that the universal frequency dependency *G*(*ω*) ∝ (−*iω*)^1/2^ of living cytoplasm also conforms to the concept of a glassy cytoplasm. Recent theoretical studies on jamming rheology have shown that this frequency dependency may be derived from the anomalous relaxation modes in the disordered medium close to the jamming transition, crowded with colloidal objects with slippery interfaces (38, 39). In a critical point scenario, the system is marginally stable; it displays excess relaxation modes at low frequencies and becomes highly susceptible to external perturbations (91). When metabolic activity was decreased in ATP-depleted cells, the elastic plateau arose as *G*(*ω*) = *G*_0_ + *A*(−*iω*)^1/2^. Similar viscoelastic behavior with an elastic plateau has been commonly observed in glassy or jammed suspensions which were quenched beyond jamming (39, 92, 93). However, the plateau *G*_0_ disappeared when they are driven out of equilibrium under the application of external flow (40), and only *G*(*ω*) = *A*(−*iω*)^1/2^ remains, much like in the living cytoplasm. The glassy suspensions exhibited the rheological properties typical for critical jamming [*i.e*., *G*(*ω*) ∝ (−*iω*)^1/2^] when they were mechanically driven out of equilibrium. Our results in cells may indicate that living cytoplasm also reflects such a critical point-like situation.

Cytoskeletons also show glassiness when they are driven out of equilibrium, *e.g*., when the network of semiflexible polymers is forced to flow beyond the linear response regime (94, 95). A glassy worm-like chain model explains such behavior by assuming that the height of the free-energy barrier for the structural modification of the network is widely spread (96). Even if thermal activation is not sufficient to overcome the barrier within the experiment duration, the broad spectrum of the energy landscape becomes apparent in the nonlinear flow behavior. At equilibrium, the linear response of *in vitro* cytoskeletons is understood as a network of semi-flexible polymers (33, 46, 97). However, cytoskeletons are driven out of equilibrium in living cells by active mechano-enzymes (81). With non-thermal perturbation, structural relaxations are induced in the cytoskeleton of living cells, leading to glassy cytoskeletal rheology. Such glassy behavior may be described by an effective noise temperature as determined in the theory of “soft glassy rheology” (22). Indeed, glassy rheology was observed in cytoskeletons at cell surfaces *in vivo* (60, 63), but in a way distinct from colloidal crowding. *G*(*ω*) of the cell cortex cytoskeleton showed widely spread power-law exponents [*β* for *G*(*ω*) ∝ (−*iω*)^*β*^], depending on the situations and the types of cells.

On the other hand, *G*(*ω*) = *A*(−*iω*)^1/2^ seems to be universally observed in the intracellular cytoplasm of MDCK cells, mouse embryonic stem cells, *etc*, as we will report elsewhere. Not only the power-law exponent (*β =* 0.5), but the prefactor *A* also does not largely differ for different kind of cells, as long as cells and the probe particles are prepared such that the intracellular cytoplasm (but not the cytoskeleton) are measured, as done in this study. Again, we emphasize that these results are consistent with glassy materials close to jamming. One way to disprove theoretical models listed above is to seek the reason (1) why this universal behavior appears specifically for cytoplasmic mechanics in living cells and (2) why normal cells do not fluidize beyond the critical jamming point, but stay there. These questions may be related to the marginal stability of active cytoplasm; the activity of mechano-enzymes is regulated in a specific manner to achieve a situation similar to critical jamming, as we will discuss in a forthcoming paper (Fig. 6) (84).

**FIGURE 6.**
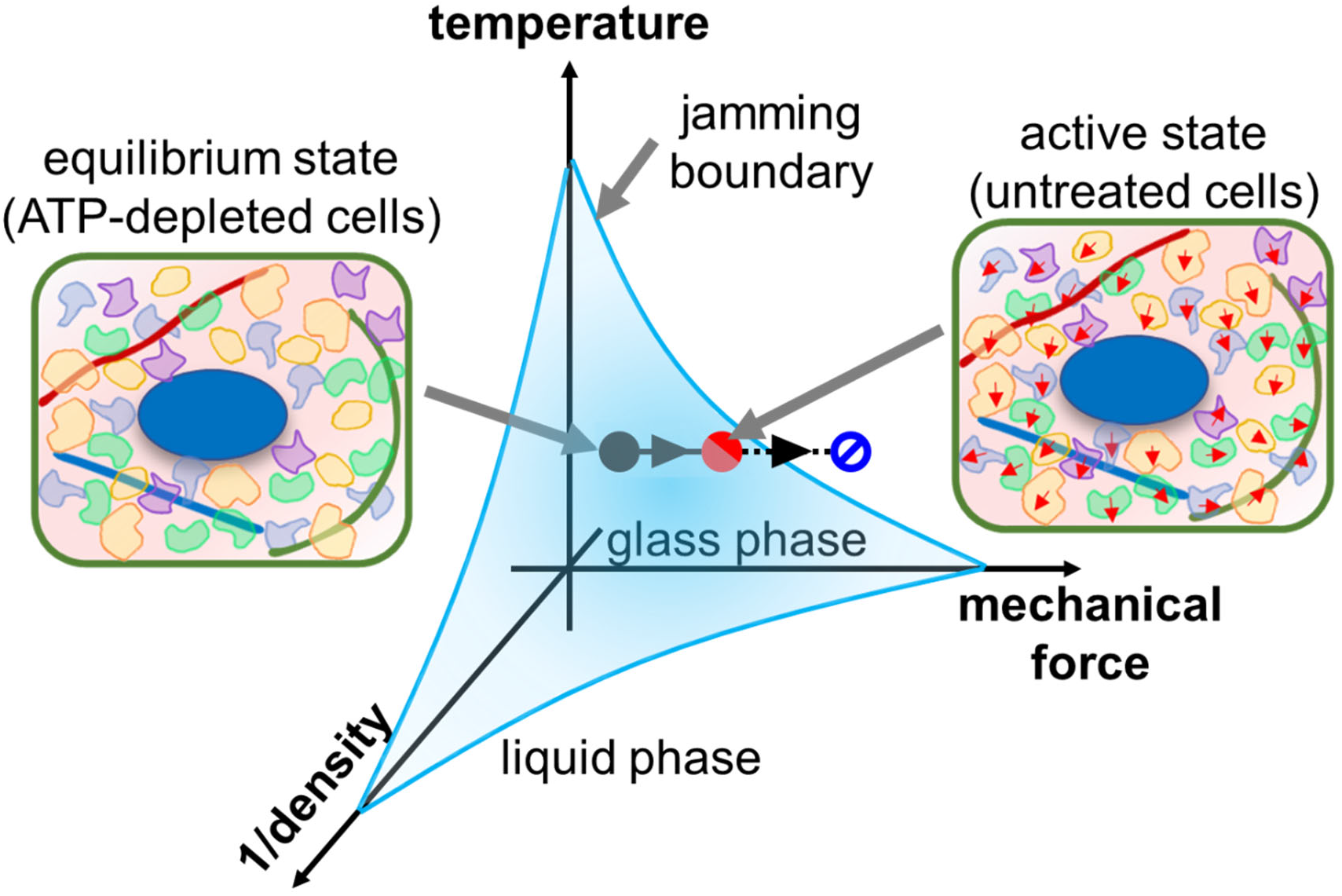
Jamming phase diagram. Energy-depleted cells are in a glassy state due to molecular crowding (black circle). The living cytoplasm, however, reaches a marginal glass phase (red circle) due to mechanical force, but does not enter a liquid phase (blue circle).

The interior of a biological cell is an enormously complex system, made up of an innumerous number of molecule types and is maintained under continuous metabolic turnover. It is therefore challenging to unequivocally prove a physical model of cells by solely investigating cells *in vivo*. Even if one inhibits or enhances a specific biochemical pathway to achieve more detailed control of metabolism, cells try to maintain homeostasis by bypassing the path in the complex network of metabolism. Thus, it would be necessary to establish a simplified model system that includes only the essences of living cytoplasm: activity and crowding. During the last decade, a model system collectively referred as “active matter” has been intensively investigated. Active matter is composed of objects which convert the internal or ambient source of energy into mechanical form. During the dissipation of the input energy, novel collective behavior of the objects was seen to emerge (98, 99). On the other hand, the physics of densely packed disordered materials (glass and jamming), especially the mechanism for dynamic arrest and melting, has been a key topic of non-equilibrium statistical mechanics over 100 years (90, 100). Recently, the active matter and glass research communities have begun to share a common interest in investigating dense active matter or active glasses (101) which we believe is a good theoretical model of living cytoplasm. Whereas fluctuations and dynamics have been investigated so far (102–108), physical and mechanical properties behind readily observable phenomena have been rarely measured. In this study, we found that mechanical properties predicted for critical jamming somehow emerge in the living cytoplasm even though some of the fundamental premises of jamming theory do not apply. Therefore, to reveal the nature of jamming rheology in the cytoplasm, simplified dense active matter which allow for control over density, active force and other conditions is desirable for future work.

## CONCLUSION

In this study, we investigated the mechanical properties of HeLa cells in a confluent epithelial monolayer. The probe particle’s displacement in response to a sinusoidal optical-trapping force was observed using a feedback-tracking microrheology technique (6, 20, 35). The complex shear modulus *G*(*ω*) of cytoplasm surrounding the probe was then obtained.

In contrast to prior measurements of the cell’s cortex, the intracellular shear viscoelastic modulus showed a frequency dependency *G*(*ω*) ∝ (−*iω*)^1/2^, which was seldom affected by the cell-cycle progression during interphase and the disruption of the actin cytoskeleton.

On the other hand, ATP depletion clearly affected the mechanical properties of the cell interior; an elastic plateau at low frequencies emerged in ATP-depleted cells. These observations are consistent with the recently emerging idea that the cell interior is glassy (3, 7). Mechano-enzymes such as molecular motors enhance structural relaxations (yielding) by actively stirring the cytoplasm, as is typical in glassy media under mechanical loads. Our results imply that the intracellular cytoplasm is fluidized such that they approach the critical jamming state, characterized by *G*(*ω*) ∝ (−*iω*)^1/2^(38, 91). Instead of complete fluidization, the slow relaxation typical of a marginally stabilized system was observed in the range of frequencies measured. We note that metabolism-dependent glassy mechanics has various implications for biology. For example, a positive feedback between mechanics and metabolism may occur in cells, as a cytoplasm fluidized by non-thermal activity may facilitate metabolic processes (5, 80, 109–112).

## ACKNOWLEDGEMENTS

This work was supported by JSPS KAKENHI Grant Number JP22H04848, JP21H01048, JP20H05536, JP20H00128. We thank Christoph Schmidt in Duke University, and Atsushi Ikeda in the University of Tokyo for helpful discussions and technical support.

## AUTHOR CONTRIBUTION

H.E., K.U., K.N., W.N., and S.I. collected/analyzed MR data. T.M. performed biochemical assays and immunocytochemistry. H.E., Y.S., and D.M. analyzed/discussed the results. D.M. designed/supervised the project, and wrote the manuscript with the support of H. E. and Y.S.

