## Supplementary for "Activity-dependent glassy cell mechanics I : Mechanical properties measured with active microrheology"

### Supplementary Materials

#### 1. *Feedback-tracking active microrheology*

We briefly provide the essence of the feedback-tracking active microrheology to help readers understand the main text. For details of the technique, see the articles given elsewhere (1, 2). We use optical trapping to apply an external force on a probe particle (3, 4), and use laser interferometry to precisely measure its position (5, 6). In order to track a probe particle that is vigorously fluctuating in a living cell, the piezo-mechanical sample stage is moved under the feedback control. As shown in Fig. S1A, the displacement of the probe particle from the probe laser $u_{\mathrm{QPD}}$, which is measured by laser interferometry, is used to control the position of the piezo stage $u_{\mathrm{stage}}$ as

$$u_{\mathrm{stage}}(t) = 1/\tau \int u_{\mathrm{QPD}} dt, \qquad (1)$$

where $\tau$ (typically set at 10 ms ~ 100 ms) is the delay time for the feedback response of our experimental setup. The displacement $u_{\mathrm{FB}}$ of the probe in the coordinate system traveling with the feedback-controlled piezo stage is given as $u_{\mathrm{FB}} = u_{\mathrm{QPD}} + u_{\mathrm{stage}}$ (Fig. 1B). From Eq. (1), the frequency-response relation is obtained as,

$$\hat{u}_{\mathrm{FB}} = (1 - i\omega\tau)\hat{u}_{\mathrm{stage}} = (1 - 1/i\omega\tau)\hat{u}_{\mathrm{QPD}}, \qquad (2)$$

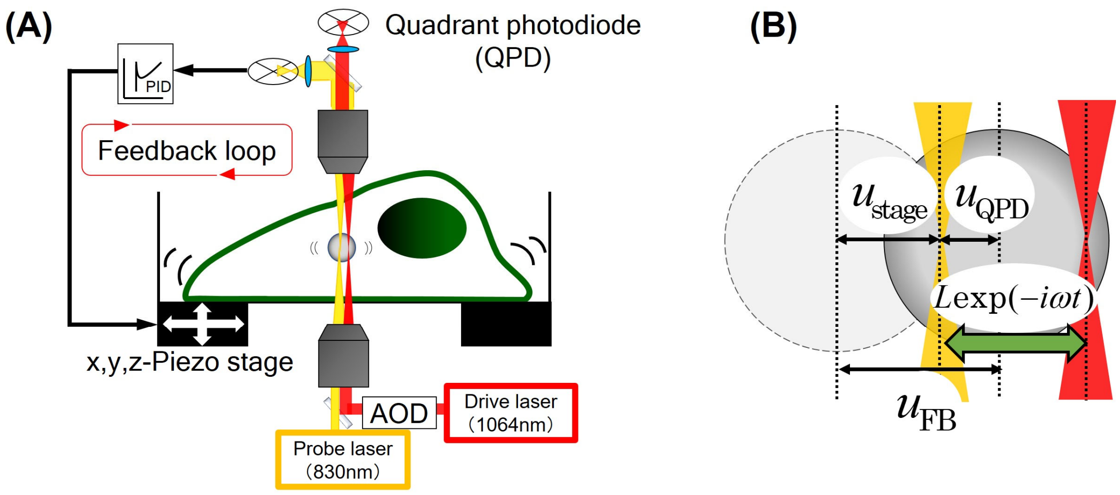


Fig. S1: (A) Schematic of the feedback-tracking microrheology setup. Fixed probe laser is used to measure $u_{\mathrm{QPD}}$ using quadrant photodiode (QPD). Drive laser is used for AMR while oscillating with acousto-optic deflector (AOD). (B) Displacements of a probe $u(t)$ is obtained by the sum of the displacement of the piezo stage $u_{\mathrm{stage}}(t)$ and the displacement from the focus of the probe laser $u_{\mathrm{QPD}}(t)$. For AMR, the focus of the drive laser is moved sinusoidally around the focus of the probe laser $L\exp(-i\omega t)$.

where $\omega$ is the angular frequency. Hereafter, ^ denotes the amplitude of the sinusoidal signal, *e.g.* $u_{\mathrm{FB}}(t)=\hat{u}_{\mathrm{FB}}(\omega)\exp(-i\omega t)$. Eq. (2) denotes that $u_{\mathrm{QPD}}$ and $u_{\mathrm{stage}}$ are high- and low-pass filtered from the total probe displacement $u_{\mathrm{FB}}(t)$. Since the slow/large displacements of the probe are tracked by piezo stage as $u_{\mathrm{stage}}$ and the fast/small displacements are measured with QPD as $u_{\mathrm{QPD}}$, the total displacement $u_{\mathrm{FB}}=u_{\mathrm{QPD}}+u_{\mathrm{stage}}$ measured by feedback-tracking technique has the advantages of high bandwidth and large dynamic range.

In the feedback-tracking AMR, a probe particle is manipulated by a sinusoidal oscillation of the overlapped drive laser whereas the probe laser is stationary (see Fig. S1b). The optical trapping force applied by the drive and probe laser is thus given as $k_{\mathrm{d}}L\exp(-i\omega t)-(k_{\mathrm{d}}+k_{\mathrm{p}})u_{\mathrm{QPD}}$, where $k_{\mathrm{d}}$, $k_{\mathrm{p}}$ refer to the trap stiffness of the drive and probe laser, respectively. $L$ is the amplitude of the drive laser oscillation. The Langevin equation for the probe particle that is under the feedback control is then written as

$$k_{\mathrm{p}}u_{\mathrm{QPD}}(t)+\int_{-\infty}^{t}\gamma(t-t')\dot{u}_{\mathrm{FB}}(t')dt'=k_{\mathrm{d}}\left(Le^{-i\omega t}-u_{\mathrm{QPD}}(t)\right)+f_{\mathrm{th}}(t)+f_{\mathrm{A}}(t) \qquad (3)$$

where $\dot{u}_{\mathrm{FB}}(t)$ denotes the velocity of the probe under the feedback control. $f_{\mathrm{th}}(t)$ and $f_{\mathrm{A}}(t)$ are the thermal and active non-thermal forces respectively, and $\gamma(t)$ is the friction function. For a stationary sinusoidal response, the time average of Eq. (3) yields the frequency-dependent response amplitude which we rewrite as,

$$\langle\hat{u}_{\mathrm{FB}}(\omega)\rangle=\langle\hat{u}_{\mathrm{QPD}}(\omega)\rangle+\langle\hat{u}_{\mathrm{sage}}(\omega)\rangle=\alpha(\omega)\left\{\hat{F}(\omega)-\left(k_{\mathrm{d}}+k_{\mathrm{p}}\right)\langle\hat{u}_{\mathrm{QPD}}(\omega)\rangle\right\}. \qquad (4)$$

Here, $\hat{F}(\omega)\equiv k_{\mathrm{d}}L$ is introduced as the "apparent" force amplitude which would be applied to the probe particle if its position is fixed. $\alpha(\omega)\equiv-1/i\omega\tilde{\gamma}$ is the intrinsic response function of a probe in the media, and ~ indicates Fourier transform. By substituting Eq. (2) into Eq. (4), we obtain

$$\alpha(\omega)=\frac{\langle\hat{u}_{\mathrm{FB}}(\omega)\rangle}{\hat{F}(\omega)-\beta\langle\hat{u}_{\mathrm{FB}}(\omega)\rangle}, \qquad (5)$$

where $\beta\equiv(k_{\mathrm{d}}+k_{\mathrm{p}})/(1-i\omega\tau)$. The complex shear viscoelastic modulus $G(\omega)$ of the surrounding material is then obtained via the Einstein-Stokes relation extended to frequency response, $G(\omega)=1/6\pi\alpha(\omega)a$ where $a$ is the radius of the probe.

#### 2. *Probe fluctuations after prolonged exposure of trapping laser*

It has been known that laser light is toxic for living cells. Although the molecular mechanism of the toxicity is largely elusive (7-9), laser irradiation reduces the motility (10) and other physiological activities of E. coli (11, 12), depending on the strength and wavelength of the laser. When conducting the optical trapping and the laser interferometry (LI), the laser is focused and therefore intensified at the probe particle. Physical properties of cytoplasm, especially in the region close to the probe particle, might be affected by the irradiation of the laser. We therefore tested whether the probe particle's movements were changed by the prolonged exposure of the laser light. A probe particle in each cell was irradiated with 1064 nm laser. After 0 min, 60 min, and 180 min of laser irradiation with the designated power, 1064 nm was turned off for a short while and PMR was quickly performed with 830 nm laser. The probe particles were stably tracked with the feedback during the laser irradiation and PMR measurements.

Fig. S2 shows the power spectral density $\left\langle |\tilde{u}(\omega)|^2 \right\rangle$ of probe fluctuations which is scaled as $\omega\left\langle |\tilde{u}(\omega)|^2 \right\rangle / 2k_B T$. The cytoplasmic fluctuation was decreased when cells were exposed to the laser for a prolonged period of time with more than a couple of mW power. It is to be noted that the optical-trapping experiments are usually performed using the laser with more than several mW intensity (13). Optical trapping with a weaker laser is not stable even in quiescent specimen at the thermodynamic equilibrium. In order to stably track a probe with such a weak trapping laser, it is necessary to use the feedback-tracking technique.

We observed that cells that showed abnormally-enhanced metabolic activity referred to as Warburg effect (14) are especially sensitive to the laser irradiation (Data not shown). It is therefore likely that the sensitivity of photochemical effects to the laser irradiation depends on the metabolic activity of cells. HeLa cells used in this study that rapidly grow in high glucose media may also show the abnormally enhanced glycolysis under aerobic condition that is the hallmark for the late-stage fully-developed cancer (15-17). Although other normal cells might be less sensitive to laser irradiation, henceforth we decided to keep the laser power less than 1 mW for conducting MR experiments in living cells.

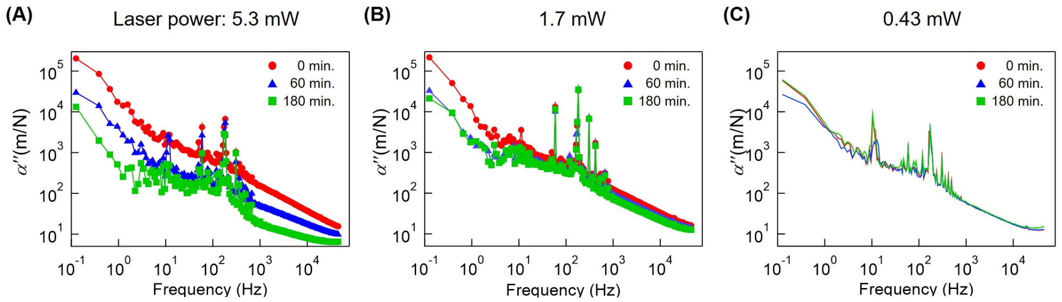


Fig. S2: Scaled power spectral densities $\omega\left\langle\left|\tilde{u}(\omega)\right|^{2}\right\rangle / 2k_{\mathrm{B}}T$ of probe fluctuations in MDCK cells. Probe particles started to be illuminated with 1064 nm laser at $t$ = 0 min.  results at $t$ = 0 min., $t$ = 60 min. $t$ = 180 min. are given in red, blue, green symbols, respectively. Experimental conditions are the same except that the laser power for illumination was 5.3 mW in (A), 1.7 mW in (B), and 0.43 mW in (C), respectively.

*3.* $G''/\omega^{0.5}$ *in untreated and actin-disrupted HeLa cells.*

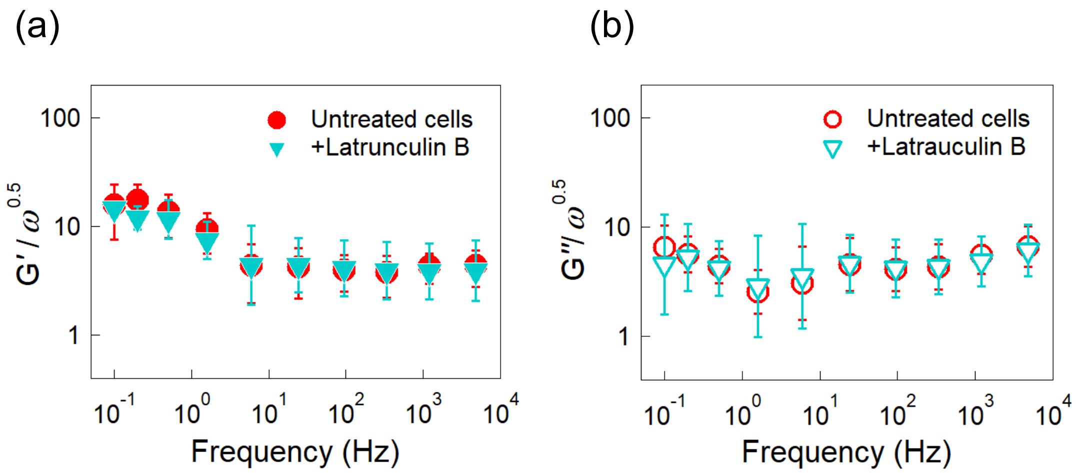


Fig. S3: (A) $G'/\omega^{0.5}$ and (B) $G''/\omega^{0.5}$ of untreated HeLa cells (circles) and treated by 10 μM Latrunculin B (inverted triangles). The same cells were measured before and after adding Latrunculin B to culture media (n = 4). Both $G'(\omega)$ and $G''(\omega)$ did not change significantly. Bars indicate the log-normal SD. In the same manner as the Fig. 3 in the main text where Cytochalasin D was used, actin disruption did not alter intracellular mechanical properties.

#### *4. $G'(\omega)$ before and after ATP depletion measured using the same probe particle in the same cells*

Intracellular mechanical properties were observed with AMR before and after imposing ATP depletion on cells. After MR experiments were performed at 37 °C in L-15 medium with 10 % FBS (w/o ATP depletion), culture medium was replaced to that for ATP depletion (L-15 medium with 50 mM 2-deoxy-D-glucose, 10 mM sodium azide, without FBS). Cells were then incubated at 37 °C on the MR setup for ~10 hours prior to the next MR measurement so that cells consume up ATP stored in cells.

$G'(\omega)$ of untreated and ATP-depleted HeLa cells are shown in Fig S4. Filled symbols and curves represent the real part *G'* of the complex shear moduli of ATP-depleted HeLa cells and those of normal untreated HeLa cells. The data shown with the same color indicates that they were measured using the same probe particle (n = 3). Note that $G'(\omega)$ always increased after ATP depletion; *G'* in the low frequency range (0.1 Hz to 1.6 Hz) is about 2 to 10 times higher in ATP-depleted HeLa cells than those in untreated HeLa cells.

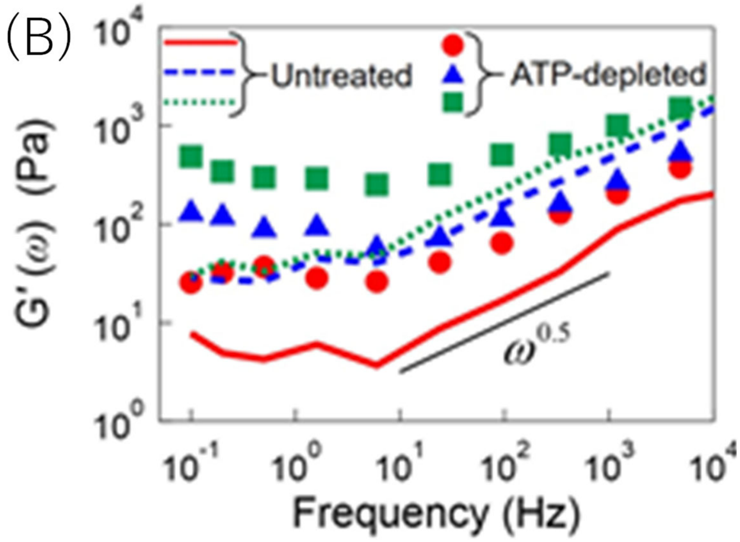


Fig. S4 $G'(\omega)$ of untreated (broken curves) and ATP-depleted HeLa cells (filled circles). The same-colored curve and circle indicate the results measured in the same cell before and after ATP depletion. In all cases, $G'(\omega)$ increased after ATP depletion especially at low frequencies.

#### 5. *Apoptotic and Necrotic Cell Detection*

A total of 2 × $10^5$ cells were plated on a poly-lysine-coated glass-bottom dish (Matsunami, D1131H) and incubated for 24 h at 37°C in 5% $CO_2$. Following incubation, the cells were incubated with either: (1) control medium [phenol-red free DMEM (Thermo Fisher, 31053028) supplemented with 4 mM glutamine (Thermo Fisher, 25030081) and 1% Penicillin-Streptomycin Solution (Sigma-Aldrich, P4333)]; (2) control medium supplemented with 50 mM 2-deoxyglucose (Nacalai, 10722-11) and 10 mM $NaN_3$ (WAKO, 199-11095); or (3) control medium supplemented with 10% fetal bovine serum (Thermo Fisher, 10270-106) and 0.5 μM staurosporine (MERCK, 569396-100UGCN), for 7 h at 37°C in 5% $CO_2$. Subsequently, the cells were collected and stained using the Cell Meter Apoptotic and Necrotic Multiplexing Detection Kit II (AAT Bioquest, 22843), according to the manufacturer's instructions. Briefly, the cells were resuspended in 200 μl of assay buffer and incubated with 2 μl of 100× Apopxin Deep Red, 1 μl of 200× Nuclear Green DCS1, and 1 μl of 200× CytoCalcein Violet 450 for 40 min at 25°C, protected from light. Following the incubation, cells were washed twice with the assay buffer and then resuspended with 200 μl of the assay buffer. The cell mixture was plated on a glass-bottom dish and then left for 5 min before imaging commenced.

For epifluorescence microscopy images, Apopxin Deep Red, Nuclear Green DCS1, and CytoCalcein Violet 450 excitation was performed using an Intensilight Mercury-Fiber Illuminator (Nikon). Data were processed through an mCherry-B-NTE-ZERO filter (Semrock), GFP-A-Basic filter (Semrock), and BFP-A-Basic filter (Semrock) for Apopxin Deep Red, Nuclear Green DCS1, and CytoCalcein Violet 450 imaging, respectively. The cells were viewed under a 40× objective (Plan Apochromat Lambda Series, Nikon) mounted on an inverted Eclipse Ti2-E microscope (Nikon) and imaged using a Zyla 4.2 PLUS sCMOS camera (Oxford Instruments). Imaging data were processed using the NIS-Elements Advanced Research imaging software (Nikon).

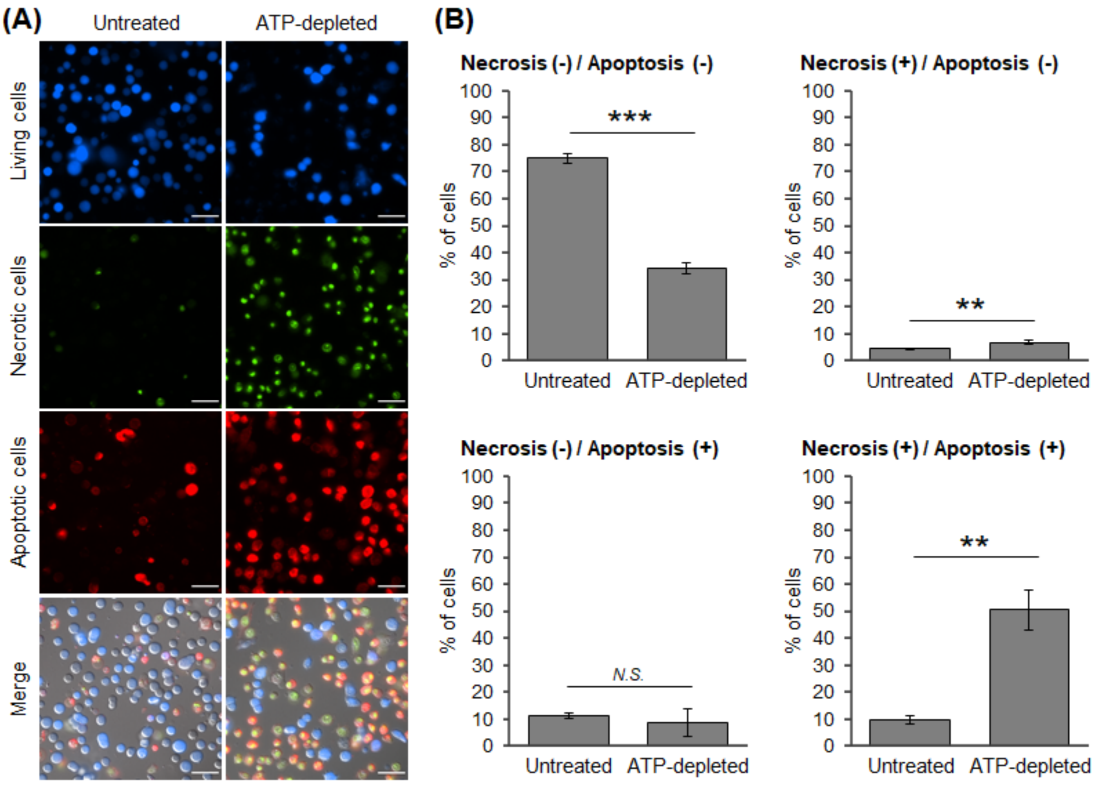


Fig. S5: HeLa cells were treated with 50 mM 2-deoxyglucose and 10 mM $NaN_3$ (ATP-depleted) for 7 h, and Apopxin Deep Red-, Nuclear Green DCS1-, and CytoCalcein Violet 450-positive cells were counted. **(A)** Representative images of collected cells from the indicated condition are shown. Scale bar = 50 μm. **(B)** Quantification was performed on three independent experiments. All data are presented as mean ± standard deviation. Total analyzed cell number: untreated cells = 4,672 (1,183 + 1,598 + 1,891) cells; ATP-depleted cells = 6,707 (2,001 + 2,188 + 2,518) cells. Living = CytoCalcein Violet 450-positive, Necrotic = Nuclear Green DCS1-positive, Apoptotic = Apopxin Deep Red-positive, Both = Nuclear Green DCS1 and Apopxin Deep Red-positive. N.S.: statistically non-significant, **: $p < 0.01$. ***: $p < 0.001$ (Student's t-test).

#### 6. *Typical experimental condition*

In this section, we listed the typical values of displacement amplitude of drive laser $A_d$, amplitude of the force form drive laser $F_d$, and displacement amplitude of aprobe particle $A_p$ in Table S1. Since $A_p$ is much smaller than the diameter of the probe particle, 1μm, we should measure linear viscoelasticity of the cytoplasm.

| $f$ (Hz) | 0.1 | 0.2 | 0.5 | 1.6 | 6 | 24 | 94 | 340 | 1200 | 4800 |
|---|---|---|---|---|---|---|---|---|---|---|
| $A_d$ (nm) | 67 | 67 | 63 | 98 | 98 | 117 | 119 | 121 | 105 | 127 |
| $F_d$ (pN) | 0.31 | 0.30 | 0.28 | 0.26 | 0.26 | 0.28 | 0.28 | 0.29 | 0.29 | 0.38 |
| $A_p$ (nm) | 5.7 | 3.6 | 2.8 | 1.2 | 0.38 | 0.28 | 0.14 | 0.078 | 0.041 | 0.022 |

Table. S1: Typical experimental condition of AMR. $f$ : frequency of drive laser. $A_d$: Displacement amplitude of drive laser. $F_d$: Amplitude of applied force from drive laser. $A_p$: Displacement amplitude of a probe particle.

#### 7. *List of P values*

Tables of P values of MR measurement in Fig. 3, 4, and 5 are provided below.

| $f$ (Hz) | 0.1 | 0.2 | 0.5 | 1.6 | 6 | 24 | 94 | 340 | 1200 | 4800 |
|---|---|---|---|---|---|---|---|---|---|---|
| *P of G'* | 1 | 2.99 x $10^{-1}$ | 5.03 x $10^{-2}$ | 5.03 x $10^{-2}$ | 1.90 x $10^{-1}$ | 7.96 x $10^{-1}$ | 3.40 x $10^{-1}$ | 8.63 x $10^{-1}$ | 7.96 x $10^{-1}$ | 9.31 x $10^{-1}$ |
| *P of G''* | 4.81 x $10^{-1}$ | 7.12 x $10^{-2}$ | 2.22 x $10^{-1}$ | 5.46 x $10^{-1}$ | 6.66 x $10^{-1}$ | 6.05 x $10^{-1}$ | 6.66 x $10^{-1}$ | 7.96 x $10^{-1}$ | 7.30 x $10^{-1}$ | 6.66 x $10^{-1}$ |

Table. S2: P values of Fig. 3 (A) (B) in main text calculated by the Mann–Whitney U test. P values are calculated for untreated and treated cells by 50 μg/ml Cytochalasin D.

| $f$ (Hz) | 0.1 | 0.2 | 0.5 | 1.6 | 6 | 24 | 94 | 340 | 1200 | 4800 |
|---|---|---|---|---|---|---|---|---|---|---|
| *P of G'* | 3.60 x $10^{-2}$ | 5.97 x $10^{-2}$ | 3.02 x $10^{-2}$ | 1.16 x $10^{-2}$ | 3.37 x $10^{-1}$ | 8.17 x $10^{-2}$ | 3.02 x $10^{-2}$ | 4.28 x $10^{-2}$ | 2.52 x $10^{-2}$ | 5.07 x $10^{-2}$ |
| *P of G''* | 1.01 x $10^{-1}$ | 4.94 x $10^{-1}$ | 3.02 x $10^{-2}$ | 2.41 x $10^{-1}$ | 5.94 x $10^{-1}$ | 2.09 x $10^{-2}$ | 2.14 x $10^{-1}$ | 6.70 x $10^{-2}$ | 2.41 x $10^{-1}$ | 3.37 x $10^{-1}$ |

Table. S3: P values of Fig. 4 (A) (B) in main text calculated by the Mann–Whitney U test. P values are calculated for cells in G1 and S/G2 phases.

| *f* (Hz) | 0.1 | 0.2 | 0.5 | 1.6 | 6 | 24 | 94 | 340 | 1200 | 4800 |
|---|---|---|---|---|---|---|---|---|---|---|
| *P of G'* | 3.24 x $10^{-4}$ | 8.28 x $10^{-5}$ | 2.59 x $10^{-4}$ | 3.50 x $10^{-4}$ | 3.89 x $10^{-3}$ | 1.28 x$10^{-2}$ | 1.24 x $10^{-1}$ | 2.98 x $10^{-1}$ | 2.22 x $10^{-1}$ | 6.21 x $10^{-1}$ |
| *P of G''* | 7.40 x $10^{-1}$ | 7.60 x $10^{-2}$ | 2.89 x $10^{-1}$ | 8.69 x $10^{-1}$ | 4.17 x $10^{-2}$ | 3.42 x $10^{-1}$ | 1 | 4.15 x $10^{-1}$ | 4.69 x $10^{-1}$ | 4.42 x $10^{-1}$ |

Table. S4: P values of Fig. 5 (C) (D) in main text calculated by the Mann–Whitney U test. P values are calculated for untreated and ATP-depleted cells.


1. Nishizawa, K., M. Bremerich, H. Ayade, C. F. Schmidt, T. Ariga, and D. Mizuno. 2017. Feedback-tracking microrheology in living cells. Sci Adv 3(9):e1700318.
2. Sugino, Y., M. Ikenaga, and D. Mizuno. 2020. Optimization of Optical Trapping and Laser Interferometry in Biological Cells. Appl Sci-Basel 10(14).
3. Mizuno, D., D. A. Head, F. C. MacKintosh, and C. F. Schmidt. 2008. Active and Passive Microrheology in Equilibrium and Nonequilibrium Systems. Macromolecules 41(19):7194-7202.
4. Hough, L. A., and H. D. Ou-Yang. 2002. Correlated motions of two hydrodynamically coupled particles confined in separate quadratic potential wells. Phys Rev E Stat Nonlin Soft Matter Phys 65(2 Pt 1):021906.
5. Gittes, F., and C. F. Schmidt. 1998. Signals and noise in micromechanical measurements. Method Cell Biol 55:129-156.
6. Gittes, F., and C. F. Schmidt. 1998. Interference model for back-focal-plane displacement detection in optical tweezers. Opt Lett 23(1):7-9.
7. Rasmussen, M. B., L. B. Oddershede, and H. Siegumfeldt. 2008. Optical tweezers cause physiological damage to Escherichia coli and Listeria bacteria. Appl Environ Microb 74(8):2441-2446.
8. Liang, H., K. T. Vu, P. Krishnan, T. C. Trang, D. Shin, S. Kimel, and M. W. Berns. 1996. Wavelength dependence of cell cloning efficiency after optical trapping. Biophys J 70(3):1529-1533.
9. Igarashi, N., S. Onoue, and Y. Tsuda. 2007. Photoreactivity of amino acids: Tryptophan-induced photochemical events via reactive oxygen species generation. Anal Sci 23(8):943-948.
10. Neuman, K. C., E. H. Chadd, G. F. Liou, K. Bergman, and S. M. Block. 1999. Characterization of photodamage to Escherichia coli in optical traps. Biophys J 77(5):2856-2863.
11. Konig, K., H. Liang, M. W. Berns, and B. J. Tromberg. 1995. Cell damage by near-IR

microbeams. Nature 377(6544):20-21.
